# The complementary roles of rare variant burden scores and common variant polygenic risk scores in genetic risk polygenic risk scores in genetic risk prediction of complex disorders

**DOI:** 10.1101/2025.09.15.676308

**Authors:** Francesco Mazzarotto, Massimo Gennarelli, Graham Murray, Emanuele F Osimo

**Author notes:** Corresponding author Emanuele F Osimo., Address: Douglas House, 18b Trumpington Rd, Cambridge, CB2 8AH, UK.

## Abstract

Polygenic risk scores (PRS) derived from common variants have improved risk prediction for complex disorders, however they capture only a fraction of heritability. One of the reasons for this is the exclusion of rare variants, despite growing evidence of their substantial contribution to genetic liability. Here we implemented the Independent Outlier Gene Count (IOGC) – a score summarising the individual burden of rare variants associated with outlying gene expression – and measured the amount of liability explained by this rare-variant score, alone and in combination with common-variant PRS, for schizophrenia, major depressive disorder, and hypertrophic cardiomyopathy in the UK Biobank. IOGC and PRS were minimally correlated (|r|<0.2), indicating the two scores capture complementary information. For schizophrenia, a model including both IOGC and PRS explained 1.13 times the adjusted liability than PRS alone. For hypertrophic cardiomyopathy, a 1.48-fold increase was observed for the combined model. People with high IOGC showed a 64% and 46% higher positive predictive value for schizophrenia and HCM, respectively, as compared to those with low IOGC. No improvement was observed for depression. These findings demonstrate that integrating rare- and common-variant scores enhances genetic risk prediction for complex conditions with a strong genetic component, supporting the inclusion of rare variant burden metrics in future genomic risk models.

## 1. Background

Historically, heritable human diseases and traits have been broadly divided into Mendelian and complex/polygenic. The former included a host of rare diseases in which, usually, a single rare variant causes a relatively conserved phenotype (Radford & Firth, 2019). Complex traits and conditions such as body height, hypertension and schizophrenia, on the other hand, are generally determined by a combined effect of genetic and environmental variables, with the genetic component constituted by several common variants, each contributing a small effect – a phenomenon known as polygenicity (Visscher et al., 2021).

Such common variants, and hence common polygenic traits, have been the object of intense scientific scrutiny – especially since the advent of inexpensive genotyping techniques – allowing the calculation of Polygenic Risk Scores (PRS) for thousands of individuals, and measuring associations of these with common traits through Genome Wide Association Studies (GWASes). GWAS power correlates largely with case-control sample size and heritability: using the example of schizophrenia, the increasing size of GWAS cohorts was paralleled by a substantial increase in the number of loci associated with the condition (Table 1).

**Table 1.** relationship between sample size and number of GWAS significant loci for schizophrenia.

| Wave | Year | Sample Size<br>(Cases + Controls) | Significant<br>Loci | Citation |
| --- | --- | --- | --- | --- |
| **PGC1** | 2011 | 9,394 + 12,462 | 7 | (The Schizophrenia Psychiatric GWAS Consortium, 2011) |
| **PGC2** | 2014 | 36,989 + 113,075 | 108 | (Pantelis et al., 2014) |
| **PGC3/**<br>**CLOZUK** | 2018 | ~40,000 + ~64,000 | 145 | (Pardiñas et al., 2018) |
| **PGC4** | 2022 | >70,000 +<br>>100,000 | 270 | (Trubetskoy et al., 2022) |

The effects of rare variants on complex conditions, on the other hand, have been more difficult to investigate due to the inherent individual rarity of these variants combined with their sheer number. The statistical techniques developed for common variants, such as GWAS, would have required case-control samples of millions of people to approach the statistical power of common-variant GWASes. As a result, rare variant association studies historically focused on highly penetrant exonic variation, mostly in the context of Mendelian conditions. However, the binary separation between rare variants-Mendelian disease and common variants-complex disease is being gradually revisited, because of several reasons:

1. Archetypical complex, highly polygenic conditions such as schizophrenia have been shown to have substantial contributions from “ultra-rare coding variants”, which have odds ratios of up to 50 (Singh et al., 2022);
2. Conditions such as hypertrophic cardiomyopathy (HCM), classically considered Mendelian, are now widely accepted as part-complex, with no single disease-causing variant found in >50% cases and with a common SNP-based heritability estimated between 17 and 29% (Mazzarotto et al., 2020)
3. More in general, many recent studies investigated the role of common variation, measured through polygenic risk scores, on rare disease penetrance (Smail et al., 2024), risk, and clinical presentation (Niemi et al., 2018), particularly in neurodevelopmental conditions.

In parallel to efforts measuring the polygenic contribution to rare disease, recent studies have started to consider the question of the contribution of rare variation to common disease. This is particularly relevant to increasing the amount of genetic variance explained by the genetic factor in conditions such as schizophrenia, where heritability is estimated in the 64-81% range (Hilker et al., 2018; Lichtenstein et al., 2009; Sullivan et al., 2003) but common SNPs only explain up to 24%, as calculated for example using SBayesS (Zeng et al., 2021).

One major hurdle in measuring the genome-wide contribution of rare variants to complex conditions is the high number of rare alleles in the population, with >99% coding variants observed in less than 1 person in 100 (Lek et al., 2016), making it challenging to distinguish risk-associated rare variants from benign bystander alleles. For this reason, new approaches are needed to efficiently prioritize categories of rare variants contributing to the risk of common polygenic conditions.

Smail et al identified >90,000 rare variants associated with outlying gene expression, collapsed them into an individual burden score called Independent Outlier Gene Count (IOGC), and demonstrated IOGC to explain a significant proportion of the liability toward obesity and related traits (Smail et al., 2022). The method is based on a development cohort with available whole genome sequencing (WGS) and transcriptomic data to select SNPs that are a) rare; b) within or around genes, but not necessarily coding; and c) associated with nearby gene outlying expression, after adjusting for common eQTLs. Burden scores are then calculated on a target population with information on disease status and genotypes, adding up the genome-wide number of genes with rare outlier-associated variants for each individual. These scores are therefore not associated with a specific condition, and therefore, unlike PRS, can be applied transversally to multiple traits or diseases.

In this work we aimed at developing an enhanced version of the IOGC burden score and quantifying:

– contribution to the total liability explained by the genetic factor;
– correlation with polygenic risk scores for each condition;
– added value to positive predictive values for each condition;

to a selected number of exemplar common conditions that had a large-sample GWAS available to calculate PRS, and selected on the basis of variable polygenicity and heritability. These included:

– schizophrenia (complex, high heritability);
– major depressive disorder (MDD, complex with lower heritability);
– HCM (part-Mendelian, part-complex).

To do so, we built upon Smail et al’s method, using GTEx to identify outlying expression-associated variants, collapsing individual carrier status into the IOGC burden score and testing its predictive performance in the UK Biobank (UKBB), with the improvement of utilising the larger and more recent release of WGS data in place of genotyping.

## 2. Methods

Please see code and data availability statements for each section below.

### 2.1 GTEx expression data: processing of RNA-seq data and outlier removal

GTEx v8 RNA sequencing data, comprising 49 tissues from 866 individuals, was processed according to methods developed by Ferraro et al. (Ferraro et al., 2020), retaining genes expressed in at least 20% individuals (TPM>0.1 and read count >6). Expression values were corrected for hidden confounders using PEER factors (Stegle et al., 2012) (15–60 depending on tissue sample size) and adjusted for the lead cis-eQTL per gene, the first three genotype principal components, and sex. Residuals were scaled and centred to generate gene-level Z scores.

Individuals exhibiting global expression outlier patterns *–* defined as having |Z|≥2 for more than 100 genes in a tissue *–* were excluded from the corrected expression matrices to minimise bias in downstream analyses.

This generated a file containing filtered, adjusted, PEER corrected, scaled and centred expression *Z* scores for all genes in each tissue-person pair.

### 2.2 GTEx: selection of outlying expression-associated variants from WGS

Variants were selected based on the method applied by Smail et al (Smail et al., 2022), with some adjustments.

In brief, WGS variants must be inside or within 10kb of coding genes, with gene boundaries defined as the furthest upstream and downstream transcription start and end site mapped to the GRCh38 reference, respectively.

Using bedtools (Quinlan & Hall, 2010) and tabix, variants that passed all quality filters and located within the region of interest for genes with outlying expression (|Z|≥2) in ≥1 tissue of ≥1 participant were extracted from GTEx WGS data.

Downstream filtering was performed with R scripts integrating bcftools (Danecek et al., 2021) and vcftools (Danecek et al., 2011) to retain variants specifically carried by individuals with outlying expression of the corresponding gene(s). In doing so:

- we included variants associated with outlying expression of multiple genes.
- in case of variants carried by a single GTEx participant, we required outlying expression of the same gene(s) to be in the same direction across tissues, allowing the presence of tissues with non-outlying expression (|Z|<2).
- in case of variants carried by multiple individuals, we required every variant carrier to have outlying expression of the same gene(s) in the same direction in at least one tissue, without requiring tissue overlap.

Frequency-based variant filtering was applied downstream, retaining variants with a MAF <1% in UKBB.

### 2.3 UKBB: Independent Outlier Gene Count computation

To calculate rare variant burden scores for each UK Biobank participant (UKBB, accessed under application 20294), we intersected the list of variants associated with outlying gene expression in GTEx with those observed in UKBB WGS data.

Variants observed in both datasets were retained, filtering out those with MAF>1% in the UKBB.

IOGC was computed for each UKBB participant, reflecting the number of genes harbouring ≥1 of these variants.

### 2.4 UKBB: Calculation of common-variant polygenic risk scores

GWASes for the three model conditions were selected on the basis of a) recency; b) sample size; c) diversity and d) variability in the phenotype explained by the GWAS.

#### GWAS selection

For schizophrenia, we selected the latest wave of the Psychiatric Genomic Consortium (PGC) Schizophrenia working group (Trubetskoy et al., 2022). For MDD, we selected the trans-ancestry genome-wide study of depression by the Major Depressive Disorder PGC working group (Adams et al., 2025). Finally, for HCM we selected the most recent GWAS on the condition (Tadros et al., 2025).

#### UKBB phenotype definitions

Schizophrenia in UKBB was defined using first occurrence of the diagnosis (Kendall et al., 2025). Lifetime MDD defined as International Classification of Diseases ICD (version 9 or 10)-coded MDD (Howard et al., 2018). Finally, HCM was defined as any ICD-coded lifetime HCM diagnosis.

#### Polygenic risk scores calculations

Polygenic risk scores for schizophrenia, MDD and HCM were calculated for each UKBB participant based on previously published methods (Choi et al., 2020). GWAS summary statistics for each condition were downloaded from public repositories, and SNPs from each GWAS were selected to overlap UKBB imputed genotypes, removing mismatching or ambiguous SNPs. QC was performed using plink and plink2 (Chang et al., 2015; Purcell et al., 2007), and included removal of samples with high heterozygosity, or with a kinship coefficient>0.125 (relatedness up to the second degree); removal of duplicated SNPs; removal of SNPs with a MAF <1%, INFO imputation score <0.8, a *p*-value<1e-15 from the Hardy-Weinberg Equilibrium Fisher’s exact, missingness>10%, minor allele count <100.

Polygenic risk scores were calculated using PRSice-2 (Choi & O’Reilly, 2019), with covariates including sex and the first 20 PCs of the genetic relationship matrix (GRM) to account for population stratification, and the `--keep-ambig --score avg` flags. Clumping was performed on a 250kb clumping window, with a default 0.1 r^2^ threshold and no *p* threshold.

### 2.5 Assessment of IOGC scores

IOGC and PRS scores were Z-standardised and left-joined to phenotype (case/control), retaining individuals flagged for inclusion in regression analyses. Individuals were then assigned to quantile bins (default N=20; ventiles), generating factor variables for IOGC and PRS quantiles.

#### Quantile-based association and predictive value analyses

To characterise risk gradients, we fitted logistic regressions of diagnosis on IOGC quantile (reference: lowest quantile) to obtain odds ratios (ORs) and 95% Wald confidence intervals. We implemented analogous models for PRS quantiles and subgroup analyses restricting to the upper IOGC tail (e.g., quantiles >18/20) to test whether PRS gradients steepen among individuals with high rare-variant burden. Results were displayed as quantile–OR plots on a log scale with *geom_pointrange* and non-overlapping labels (ggrepel). To assess clinical utility, we estimated positive predictive values (PPVs) across pre-specified PRS centile thresholds (<15th, <25th, <50th, <75th, >95th) separately within low-vs high-IOGC strata (dichotomised at the pre-specified IOGC quantile cutoff).

#### Coefficient of determination (liability-scale, ascertainment-corrected)

The coefficient of determination (CoD) was calculated to quantify the variance explained by the genetic factor on the liability scale, after correcting for case–control ascertainment (Lee et al., 2012). Operationally, as previously done (Osimo, 2023), we:

i. fitted linear probability models of diagnosis on genetic predictors – IOGC quantile (categorical), PRS, and additive combinations thereof (e.g., PRS + IOGC-quantile) – to obtain the observed r^2^;
ii. bootstrapped r^2^ with R = 50 resamples using the “boot” package with parallel back-ends to derive standard errors (SEs) and percentile intervals; and
iii. transformed the observed r^2^ to the liability scale using the `cc_trf` function from the “r2redux” package (Momin et al., 2023), providing the population prevalence KK (1% for schizophrenia, 10% for depression, 0.2% for HCM) and the sample case fraction PP, yielding r_l_^2^ and its SE. We report 95% CIs as r_l_^2^±1.96×SE and present results as point–range plots.

## 3. Results

### 3.1 IOGC calculation and application to UKBB

GTEx gene expression data pre-processing led to the identification of 33 global outlier individuals, which were removed from further analysis. In the remaining 833 samples, 1,592 genes had outlying expression (|Z|>2) in ≥1 tissue of ≥1 individual.

Analysed GTEx WGS data comprised a total of 69.8 million sequence variants. Of the 36 million inside or within 10kb of genes we identified 315,381 (0.88%) as outlying-expression associated, with 258,251 of these (81.9%) present also in UKBB WGS with a MAF<1% and subsequently used to compute IOGC burden scores.

UKBB participants carried a median±IQR of 84±18 (mean±SD: 89±30) outlying expression-associated rare variants (range: 0-761). In computing the IOGC burden score, multiple variants in or around the same gene were collapsed and counted as one when carried by the same individual. The median IOGC burden score – hence corresponding to the median number of genes harbouring ≥1 variant of interest – in UKBB was 68±12 (mean±SD: 69±14; range: 0-243).

### 3.2 Rare variant contribution to the genetic risk of schizophrenia

The analysis for schizophrenia was conducted on a set of 440,566 UKBB participants retained by quality control steps. Of these, 1,169 were classed as people with schizophrenia according to the methods above.

Individual IOGC burden scores were significantly higher in people with schizophrenia, as compared to controls (mean±SD: 71.7±18.3 vs 69.4±14.4; Bonferroni-corrected one-tailed Mann-Whitney U test p=7.4E-3).

#### 3.2.1 Risk scores and odds ratios for schizophrenia

The OR for schizophrenia is known to vary log-linearly with quantiles of common-variant PRS for the condition, and this was the case in UKBB. *Supplementary Figure 1* shows that the risk for the top ventile of schizophrenia PRS was associated with an OR for schizophrenia of 13.6 (95% CI: 8.1-23) as compared to the lowest, with intermediate and increasing values for quantiles in between.

*Supplementary Figure 2*, on the other hand, shows that the OR for schizophrenia did not vary linearly with ventiles of IOGC, showing a significant increase only for the top two. The top ventile was associated with an OR for schizophrenia of 2.3 (1.5-3.7) as compared to the lowest.

#### 3.2.2 IOGC improves schizophrenia prediction

To measure whether rare-variant IOGC could improve risk predictions for schizophrenia, we first investigated the correlation between this score and common-variant PRS for the condition and found that to be negligible (Pearson’s correlation coefficient (PCC)=0.097; 0.095-0.1).

We then moved on to calculating the proportion of the variance for schizophrenia explained by the genetic factor on the liability scale corrected for ascertainment, or coefficient of determination (CoD) – Figure 1 shows the CoD for IOGC was 0.56% (0.05-1.06%). Conversely, common-variant schizophrenia PRS had a CoD of 6.34% (5.08-7.60%). The combination of the two rare- and common-variant scores yielded a CoD of 7.17% (5.83-8.51%), with a relative improvement on common-variant PRS alone of ∼13.1%.

**Figure 1:**
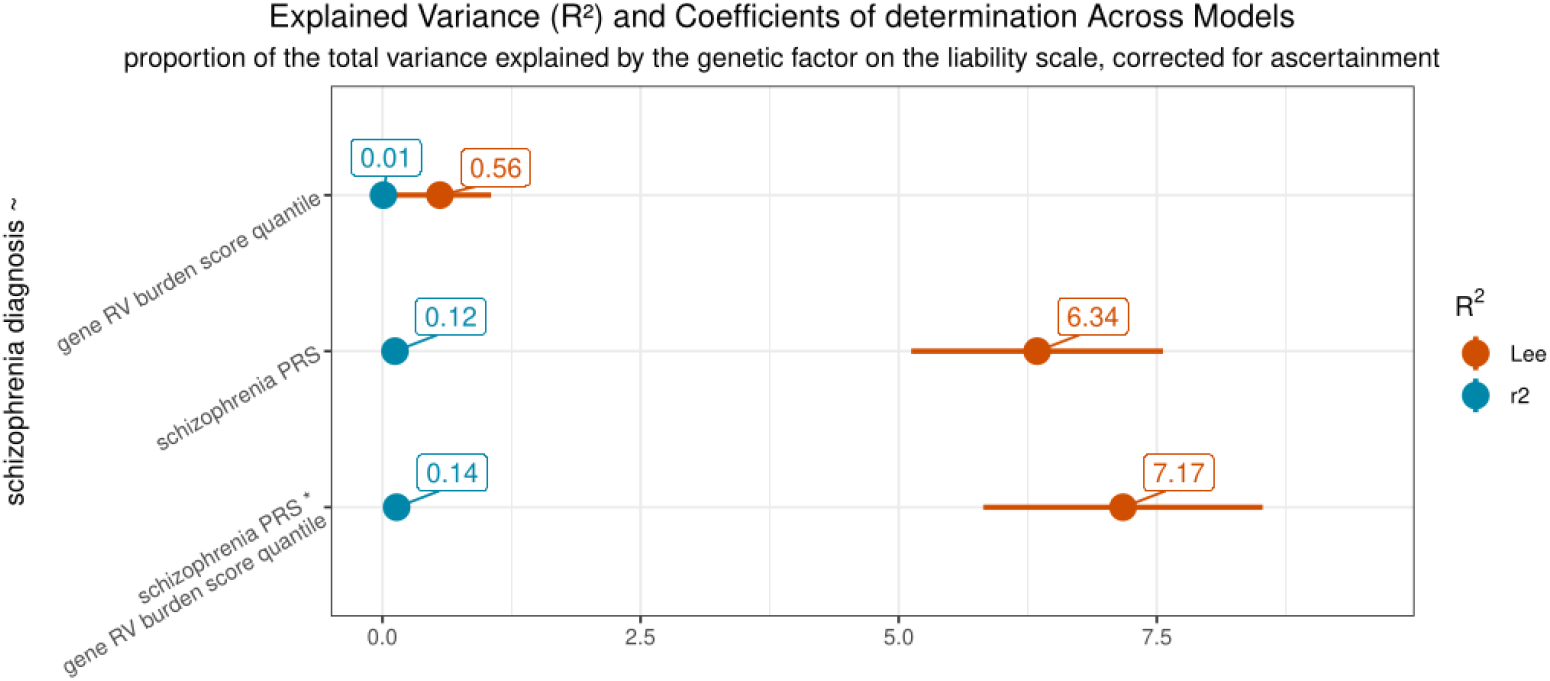
Schizophrenia variance explained by different models. The figure describes the proportion of the variance of schizophrenia explained by the genetic factor for each of three genomic scores – IOGC rare variant burden score, the original GWAS, and a combination of the two in the UKBB. The plot shows the overall CoD % and 95% confidence interval; in brick red, values are on the liability scale, corrected for ascertainment as per Lee et al., 2012, utilising the formula by Choi and O’Reilly, 2019 – or coefficients of determination (CoD). In blue the original Nagelkerke’s R2 for comparison.

PPVs for schizophrenia were calculated using common-variant PRS centiles, and dichotomising IOGC scores into low (ventiles 1-18) or high (>18), as shown in *Supplementary Figure 3*.

A high (vs low) IOGC score increased the PPV for schizophrenia for a person with any PRS score 1.64 times, from 0.25 to 0.41. A high (vs low) IOGC score also increased the PPV for schizophrenia of an individual with a PRS >95^th^ centile from 0.89 to 1.13.

### 3.3 Rare variant contribution to the genetic risk of major depressive disorder

A total of 440,631 individuals passed QC in UKBB for the MDD analysis. Of these, 26,626 were classed as people with ICD-coded lifetime MDD, according to the methods above.

Individual IOGC burden scores were not significantly different between people with MDD and controls (69.4±14.5 vs 69.4±14.4, nominal p=0.31).

#### 3.3.1 Risk scores and odds ratios for MDD

OR for MDD also varied pseudo-log-linearly with quantiles of common-variant PRS for the condition, as shown in *Supplementary Figure 4*. The risk for the top ventile of MDD PRS was associated with an OR for MDD of 11.2 (10.1-12.5) as compared to the lowest, with intermediate and increasing values for other quantiles.

Conversely, the OR for MDD did not show significant differences across IOGC quantiles (*Supplementary Figure 5*).

#### 3.3.2 Risk scores and MDD prediction

Similarly to schizophrenia, the IOGC burden score and PRS for MDD were not strongly correlated (PCC=0.08; 0.077-0.083).

Figure 2 shows the CoD of IOGC for MDD to be 0.02% (0.00-0.04%), with that for common-variant MDD PRS being 9.66% (9.28-10.05%). The combination of the two yielded a CoD of 9.80% (9.41-10.20%).

**Figure 2:**
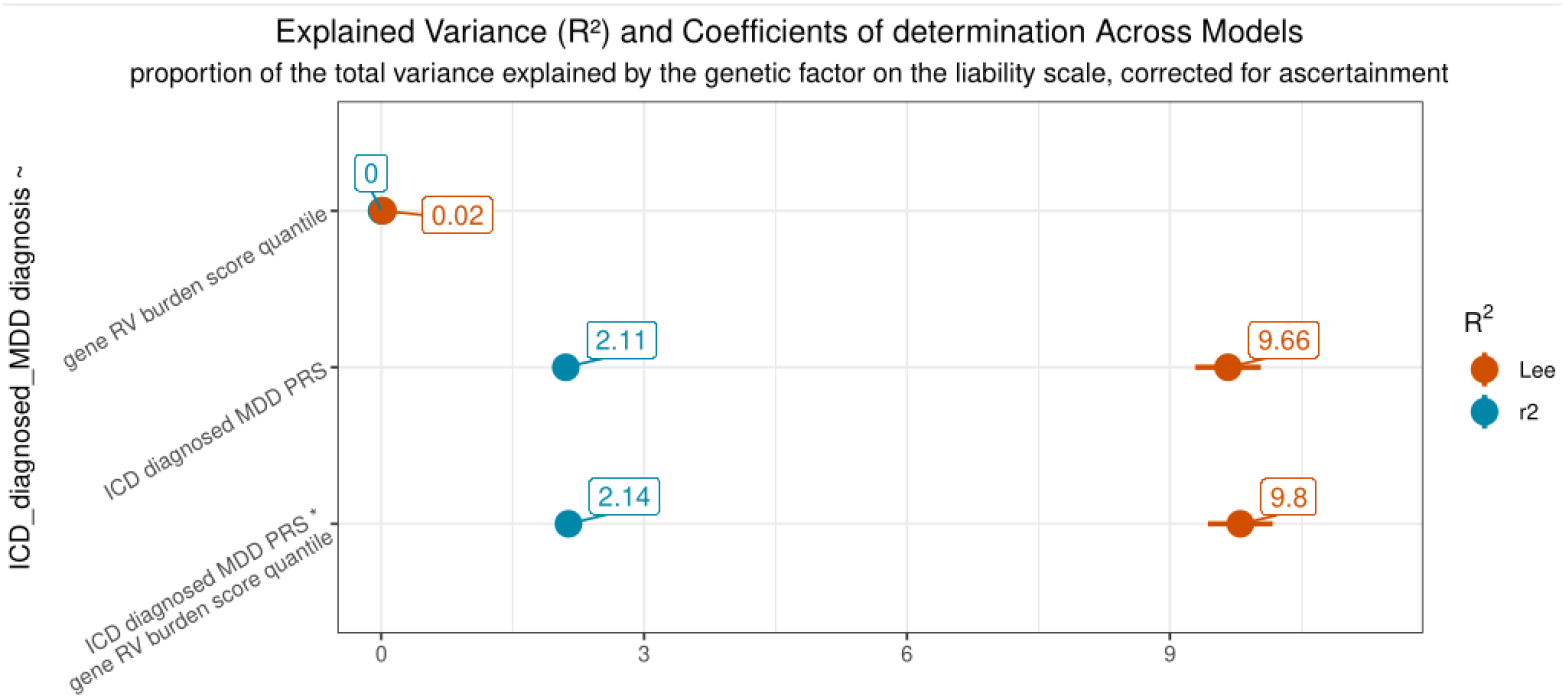
MDD variance explained by different models. The figure describes the proportion of the variance of MDD explained by the genetic factor for each of three genomic scores – IOGC rare variant burden score, the original GWAS, and a combination of the two in the UKBB. The plot shows the overall CoD % and 95% confidence interval; in brick red, values are on the liability scale, corrected for ascertainment as per Lee et al., 2012, utilising the formula by Choi and O’Reilly, 2019 – or coefficients of determination (CoD). In baby blue the original Nagelkerke’s R2 for comparison.

As done for schizophrenia, PPVs for MDD were calculated using common-variant PRS centiles, and dichotomising IOGC scores into low (ventiles 1-18) or high (>18).

A high (vs low) IOGC score showed only a very slight effect on the PPV for MDD for a person with any PRS score, from 6.03 to 6.13. For those with a PRS for MDD above the 95^th^ centile, IOGC instead decreased the PPV from 17.12 to 14.31 (*Supplementary Figure 6*).

### 3.4 Rare variant contribution to the risk of hypertrophic cardiomyopathy

440,631 individuals passed QC in UKBB for this analysis. Of these, 602 were classed as having HCM according to the methods above.

Individual IOGC scores in people with HCM appeared comparable with those of unaffected subjects (71.0±17.3 vs 69.4±14.4, nominal p=0.42).

#### 3.4.1 Risk scores and odds ratios for HCM

ORs for HCM varied with quantiles of common-variant PRS for the condition, as shown in *Supplementary Figure 7*. The risk for the top ventile of HCM PRS was associated with an OR for HCM of 10.8 (5.5-21.4) as compared to the lowest.

As observed for schizophrenia and MDD, the OR for HCM did not vary linearly with IOGC quantiles, but the top ventile was characterized by a significantly increased OR of 2.4 (1.3-4.6) compared with the bottom one (*Supplementary Figure 8*).

#### 3.4.2 Risk scores and HCM prediction

The IOGC burden score was negatively and more strongly correlated with HCM PRS – compared with PRS for schizophrenia and MDD – in UKBB (PCC = −0.178; −0.181 – −0.175).

The CoD for HCM given by the IOGC burden score and by the HCM PRS alone were 0.55% (0.13-0.96%) and 4.47% (3.04-5.90%), respectively. The combination of IOGC and HCM PRS yielded a CoD of 6.61% (4.18-9.03%), with a 47.9% improvement on PRS alone (Figure 3).

**Figure 3:**
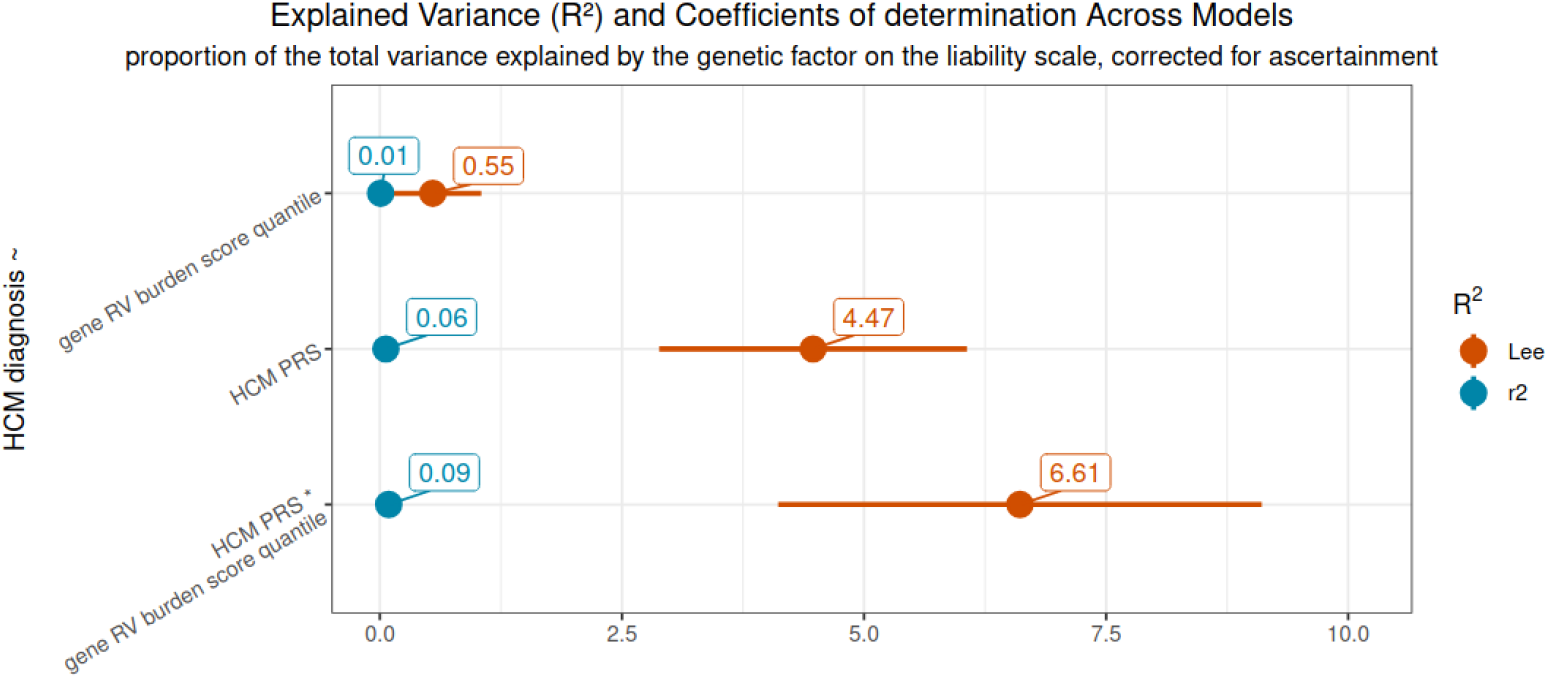
HCM variance explained by different models. The figure describes the proportion of the variance of HCM explained by the genetic factor for each of three genomic scores – IOGC rare variant burden score, the original GWAS, and a combination of the two in the UKBB. The plot shows the overall CoD % and 95% confidence interval; in brick red, values are on the liability scale, corrected for ascertainment as per Lee et al., 2012, utilising the formula by Choi and O’Reilly, 2019 – or coefficients of determination (CoD). In baby blue the original Nagelkerke’s R2 for comparison.

Dichotomised high/low IOGC generally increased PPV for HCM across all tested PRS thresholds (*Supplementary Figure 9*), with a ∼105% improvement from 0.42 to 0.86 among subjects with a PRS >95^th^ centile.

### 3.5 Sensitivity analysis with random IOGC scores

A negative control analysis was performed, after randomly swapping existing burden scores between UKBB participants thus maintaining the same distribution but randomising it across the population.

For all conditions, the CoD gains were reduced (schizophrenia PRS^*^IOGC CoD=6.86, 95%CI 5.41-7.87; MDD PRS^*^IOGC CoD=9.7, 95%CI 9.33-10.08; HCM PRS^*^IOGC CoD=5.12, 95%CI 2.71-7.52).

Crucially, PPVs for the conditions – which had shown gains in the high-vs-low IOGC comparisons for both schizophrenia and HCM – were completely reversed in this sensitivity analysis, with PPVs showing lower values when considering high-vs-low randomised IOGCs (*Supplementary figures 10-11*).

## 4. Discussion and Conclusion

### 4.1 Discussion

The disease-agnostic IOGC rare-variant burden score provided a meaningful contribution to the amount of adjusted variance explained by the genetic factor for schizophrenia and HCM, while it yielded no contribution for MDD. Individuals with an IOGC burden score above the 95th percentile were characterized by significantly increased ORs for the two former conditions, in the range between 2.3-2.4 vs the lowest 5%.

The IOGC burden score alone explained a smaller proportion of the variance explained by the genetic factor on the liability scale corrected for ascertainment (CoD) for schizophrenia and HCM (∼0.55%), compared with PRS (6.43% for schizophrenia and 4.47% for HCM), but a model combining the two scores increased the CoD by 13% and 48%, respectively. The additive – and synergistic, in the case of HCM – effects of PRS and IOGC on risk suggest the two scores capture different portions of genetic liability, as confirmed by the scarce correlation between the two scores within the same UKBB cohort (0.097 for schizophrenia and −0.178 for HCM).

Further, the dichotomisation of IOGC scores showed PPVs for each condition to improve 1.64 and 1.46-fold in schizophrenia and HCM, respectively, in subjects with a high vs low IOGC, regardless of PRS. Notably, the PPV improvement was 2.05-fold in HCM among individuals with a PRS >95^th^ centile. PPVs – on the other hand – showed a decrease when considering randomised (negative control) IOGS scores – confirming the validity of the positive results.

In its first application, IOGC scores were applied to UKBB relying on imputed genotypes (Smail et al., 2022); at the time, it was found that subjects in the top IOGC decile had a 20.8% higher risk of obesity, 62.3% higher risk of severe obesity, and underwent bariatric surgery 5.3 years earlier compared to those in the bottom decile, even after adjusting for common variant-based PRS. In this work we were able to apply IOGC scores calculated on WGS within GTEx to sequenced UKBB genomes, thus increasing the number of variants from ∼96 million (imputed) to ∼1.29 billion SNPs (WGS). As a result, IOGC scores now encompass >258,000 rare variants of potential functional relevance, compared with ∼90,000 in the previous iteration.

Schizophrenia is a common disease with a worldwide prevalence of ∼0.7% (Solmi et al., 2023), characterized by a heritability of 64-80% (Hilker et al., 2018; Lichtenstein et al., 2009; Sullivan et al., 2003) and a highly polygenic structure (Smeland et al., 2020). This means that genetic susceptibility to schizophrenia is largely due to a high number of genetic variants, each carrying a small risk. While the scientific community is beginning to study the contribution of rare genetic variation to schizophrenia risk, for the time being this effort mostly focused on ultra-rare coding variants increasing the odds of the condition manifold (Singh et al., 2022).

Our findings in the context of schizophrenia support an important role for rare variants (e.g. variants with a MAF between 0.1% and 1%), which are overlooked by both GWAS and ultra-rare variant-focused approaches due to their inherent limitations.

The same holds true for HCM, despite its lower prevalence (∼1 in 500) and radically different genetic architecture. HCM is characterized by a strong Mendelian component and, in contrast with schizophrenia, research into its genetic background historically focused mostly on rare, coding variants. To date, 18 genes are robustly associated with dominant or recessive forms of isolated, non-syndromic HCM (Hespe et al., 2025). However, in ∼50% of patients with HCM undergoing clinical genetic testing, a causative Mendelian variant in these genes is still not found (Mazzarotto et al., 2020). Different models for HCM beyond purely Mendelian inheritance (e.g. oligogenic and complex forms of HCM) are increasingly believed to explain a substantial portion of these cases, catalysing GWASes and studies on non-Mendelian variants in recent years. These studies estimated common variant-based heritability at 16% and 29% in HCM patients with vs without a known Mendelian variant, respectively, and demonstrated a contribution to HCM of intermediate-effect variants with a MAF <1% in Mendelian HCM genes (García Hernandez et al., 2025; Tadros et al., 2025). In this context, results obtained here suggest how rare variants accounted for by IOGC may explain a portion of the liability neither captured by single Mendelian HCM-causing variants nor by HCM PRS, and could contribute to shed light on intermediate-effect variants in a genome-wide fashion.

On the other hand, IOGC scores showed no effects in terms of proportion of explained genetic liability or PPV in the context of MDD. MDD is characterized by a higher lifetime prevalence (∼15%)(Adams et al., 2025), a moderate total heritability (∼37%)(Polderman et al., 2015), and by the lack of associations with rare variants with high effect sizes. Of note, MDD is characterized by a common SNP-based heritability of 8.4%, approximately three times lower than both schizophrenia and HCM (Adams et al., 2025). Overall, this relatively limited genetic contribution to MDD may represent one of the reasons why the IOGC does not seem to capture liability for this condition. In addition, MDD is a clinically heterogeneous disease; while this clinical heterogeneity is shared with schizophrenia, the substantially higher heritability of the latter may partly explain our results. A final potential explanation for the difference may lie in the putatively different pathophysiology of schizophrenia and MDD – where most hypotheses for schizophrenia seem to converge on early neurodevelopment, and altered synaptic function (Birnbaum & Weinberger, 2017). High burden scores, putatively associated with disrupted gene expression, may represent one of the causes for altered neurodevelopment in schizophrenia. In any case, this work does not provide for any indication on what the potential pathophysiology may be for such differences.

### 4.2 Limitations

This study has several limitations. First, the reliance on GTEx to identify outlying expression-associated variants represents a ceiling on the number of rare outlying variants that IOGC can identify (Ferraro et al., 2020). However, to our knowledge, this is currently the largest sample where multi-tissue RNA sequencing is available in the same sample as WGS.

Second, while the approach we applied expands into variants in non-coding regions, it is inherently limited to variants in or within 10Kb of genes (i.e. 49.9% of the genome) and with a statistically significant effect on gene expression – rather than wet lab-verified.

Third, in this work, for a given gene, we considered outlying expression in any tissue as an inclusion criterion for variants within or around the gene (Smail et al., 2022). While this is instrumental to prioritizing a higher number of variants and maximizing power, it entails accounting for transcriptional effects in tissues not necessarily relevant to the conditions of interest. However, the current methods require multiple tissue data for accurate detection.

Finally, UKBB is characterized by a “healthy volunteer” selection bias (Fry et al., 2017), including a lower disease prevalence and a generally milder disease severity in UKBB compared with the general population; further, cases for conditions including schizophrenia have been shown to include substantial differences from clinical populations (Legge et al., 2024). This will have genetic implications.

These limitations collectively warrant further developments of the IOGC approach, building upon larger transcriptomic datasets and case numbers as well as tissue-, gene- and variant-based refinements.

### 4.3 Conclusions

In this work we highlight how – pending validation in additional, larger transcriptomic datasets and case cohorts – combining rare- and common-variant genetic risk scores can yield benefits in genetic risk prediction, which may have in future clinical implications, at least for some conditions characterised by strong genetic heritability, such as schizophrenia and HCM.

Future work should focus on applying the method to more conditions and refining rare variant burden scores, e.g. by tailoring them to disease-relevant tissues, as well as extending their breadth beyond alleles with a potential regulatory effect.

## Data and code availability

While some of the data in our code is public access, most datasets require specific applications, and the data is therefore not publishable. We are however making all of our code open for collaboration and reproducibility purposes.

Unless otherwise specified, data wrangling was done in R (R Core Team, 2022) with data.table/dplyr (Wickham et al., 2017), and data were visualised using ggplot2 (Wickham, 2009).

Most scripts were ran on the University of Cambridge HPC using bash scripts.

### Processing of GTEx RNA-seq Data and Outlier Removal

GTEx data used for the analyses described in this manuscript were obtained from dbGaP Accession phs000424.v8.p2 following NIH Data Access Committee approval for project #39025.

The code for this part of the study can be accessed at https://github.com/emosyne/iogc_scz/blob/tidy/scripts/1_GTEx_data_preprocessing_and_PEER/eoutliers_calc.sh

This code builds on work by (Ferraro et al., 2020) and adapts scripts from https://github.com/nmferraro5/gtex_v8_rare_eoutliers/blob/716d43303dd31fae20869d4957b631393fb409c5/correction.md

(PEER correction), and from https://github.com/nmferraro5/gtex_v8_rare_eoutliers/blob/716d43303dd31fae20869d4957b631393fb409c5/outlier_calling.md

(outlier calling) – both under CC0 1.0 Universal license (last checked 4/9/2025). GTEx: selection of outlying expression-associated variants from WGS

Scripts to extract outlying-expression associated variants in the target regions from GTEx are available in folder https://github.com/emosyne/iogc_scz/tree/tidy/scripts/2_variant_identification_filtering_RVBS_calculation and are numbered sequentially. Scripts that are not numbered (e.g. filter_by_MAF.R) don’t need to be called by the user, as these are called by the main scripts below:

– 1_create_gene_body_bed.R was used to create a BED file with the start and end coordinates ±10Kb of all genes identified as having outlying expression in the previous steps.
– 2_extract_gtex_genebody_variants.sh extracts all GTEx PASS variants from the target regions.
– R scripts with prefix “3a” inside 3_outl_assoc_vars_ext_anytissue perform specific extraction of variants associated with outlying gene expression in GTEx participants, across 29 gene batches, applying filtering criteria described in section 2.2. The R script with prefix “3b” performs batch concatenation and further filtering.

### UKBB: Independent Outlier Gene Count computation

The sequence of scripts goes on to extract variants at relevant positions from the UKBB and compute IOGC for all participants. Extraction of variants at a specific position from UKBB requires knowledge on which WGS “chunk” contains the position of interest. Scripts to create a table with start and end coordinates of each UKBB WGS chunk are available in GitHub repository ukb_wgs_mapping_500k.

As specified in the previous section, scripts that are not numbered don’t need to be run by the user.

– 4_extract_ukb_vars.sh extracts all variants from the UKBB by genomic coordinate, using the dx-toolkit, and 5_download_ukb_vars.sh downloads extracted VCFs.
– Scripts inside 6_process_ukb_vars filter and process extracted UKBB variants. Script 6a_rm_novars_batches.sh removes empty VCFs (i.e. variants observed in GTEx but absent or non-PASS in UKBB). Scripts with the “6b” prefix sort, PASS-filter and index files in batches, splitting multi-allelic sites. Script 6c_concat_match_final_vars.sh concatenates all QCed VCFs by chromosome and retains only alleles observed in GTEx. Script 6d_create_final_file.sh concatenates chromosome files to create a final VCF.
– Scripts inside 7_tableize_final_variant_set convert the final VCFs into tables, in chromosome batches.

Script 8_create_final_tables_compute_scores.sh filters variants by their MAF in UKBB and computes IOGC burden scores for all UKBB participants, saving them in a table.

### UKBB: Calculation of common-variant polygenic risk scores

#### UKBB phenotype definitions

MDD phenotypes were build adapting code from (Howard et al., 2018), found at https://datashare.ed.ac.uk/handle/10283/3083.

#### Polygenic risk scores calculations

The steps described in the methods section above are implemented in this section of the repository: https://github.com/emosyne/iogc_scz/tree/tidy/scripts/3_PRS_score_calculation_and_plotting in steps 1-6.

#### Assessment of IOGC scores

All analyses were conducted in R – and the code can be found at https://github.com/emosyne/iogc_scz/blob/tidy/scripts/3_PRS_score_calculation_and_plotting/7_decile_and_CoD_plots.Rmd

## Acknowledgements

All research at the Department of Psychiatry in the University of Cambridge is supported by the NIHR Cambridge Biomedical Research Centre (NIHR203312) and the NIHR Applied Research Collaboration East of England. The views expressed are those of the author(s) and not necessarily those of the NIHR or the Department of Health and Social Care.

The Genotype-Tissue Expression (GTEx) Project was supported by the Common Fund of the Office of the Director of the National Institutes of Health, and by NCI, NHGRI, NHLBI, NIDA, NIMH, and NINDS.

This research has been conducted using the UK Biobank Resource under application number 20904.

## Notes

### Competing Interest Statement

The authors have declared no competing interest.

