## Supplementary Figures for "The complementary roles of rare variant burden scores and common variant polygenic risk scores in genetic risk polygenic risk scores in genetic risk prediction of complex disorders"

#### *Supplementary Figure 1: OR for Schizophrenia by Common Variant Schizophrenia PRS Quantiles*

*The quantile plot shows the odds ratios and 95% confidence intervals for schizophrenia for 20 quantiles of original GWAS PRS, from 1 (reference, and lowest PRS) to 20, in UKBB.*

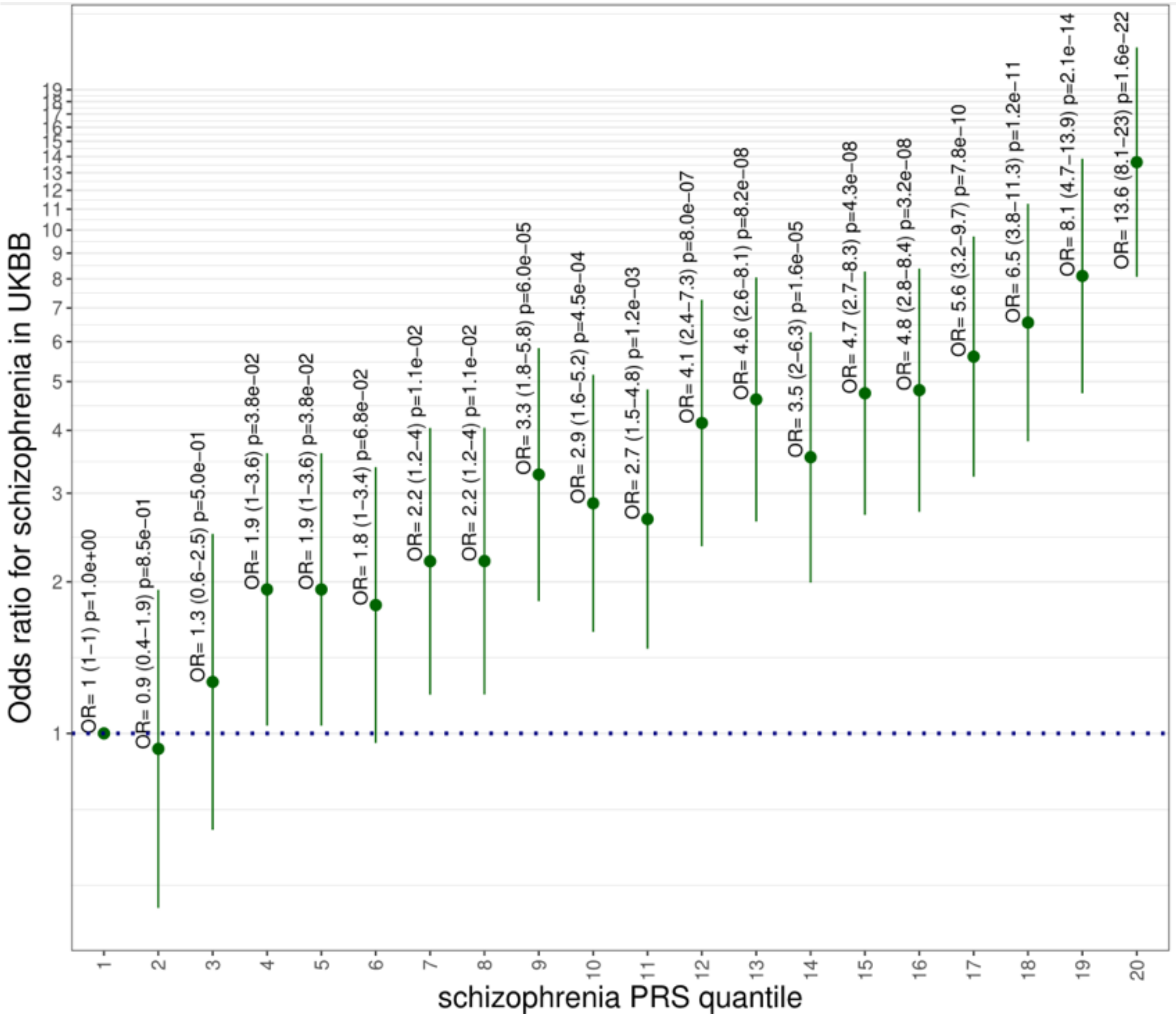

### *Supplementary Figure 2: OR for Schizophrenia by Rare Variant IOGC Quantiles*

*The quantile plot shows the odds ratios and 95% confidence intervals for schizophrenia for 20 quantiles of IOGC rare variant burden scores, from 1 (reference, and lowest PRS) to 20, in UKBB.*

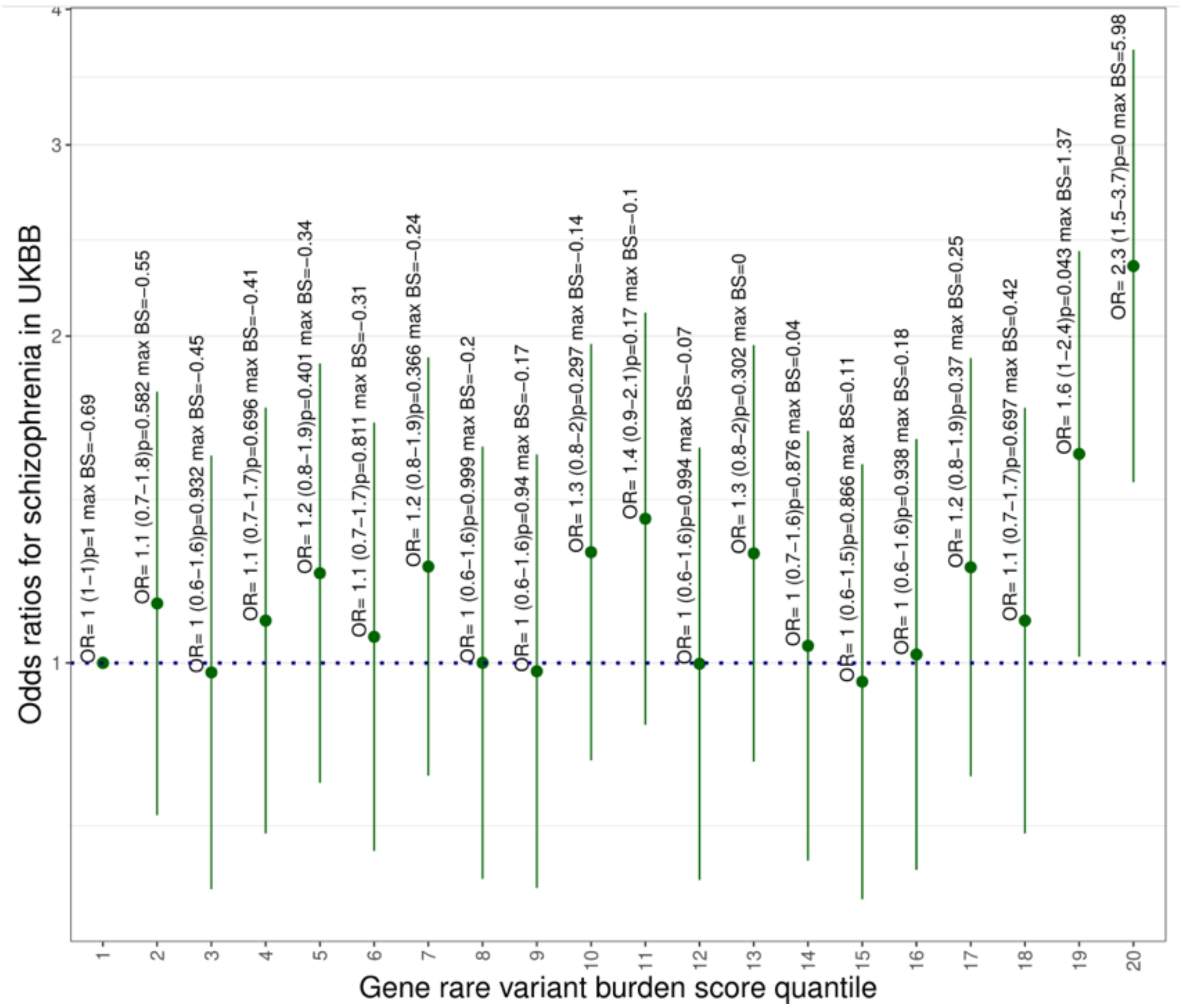

*Supplementary Figure 3: Positive Predictive Value for Schizophrenia by common-variant PRS and by high/low IOGC*

*The figure shows positive predictive values (PPVs) for schizophrenia across pre-specified PRS centile thresholds (whole sample, <15th, <25th, <50th, <75th, >95th) separately within low- vs high-IOGC strata (dichotomised at the pre-specified IOGC quantile cutoff): in blue, low IOGC; in red, high IOGC.*

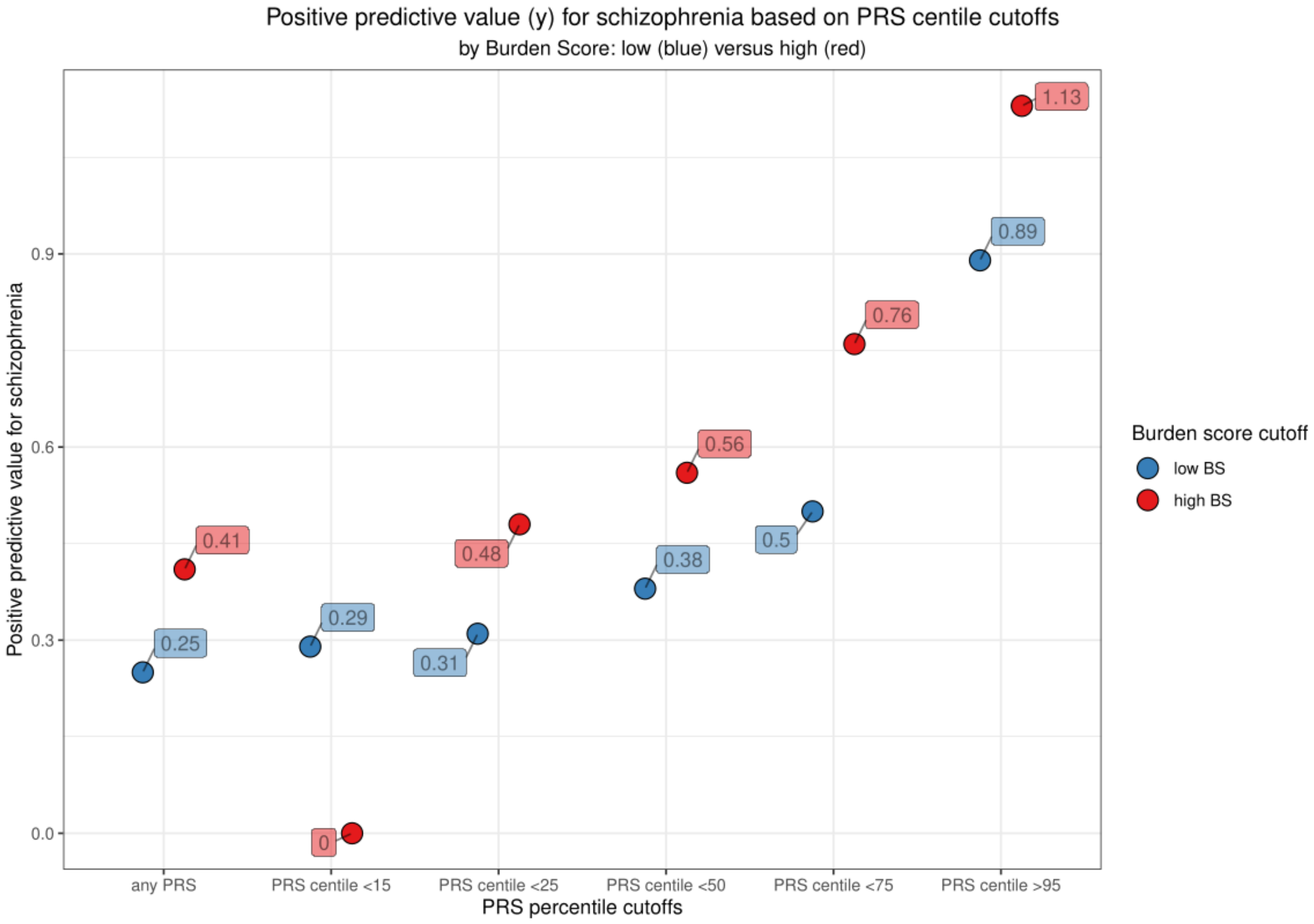

*Supplementary Figure 4: OR for MDD by Common Variant MDD PRS Quantiles*

*The quantile plot shows the odds ratios and 95% confidence intervals for MDD for 20 quantiles of original GWAS PRS, from 1 (reference, and lowest PRS) to 20, in UKBB.*

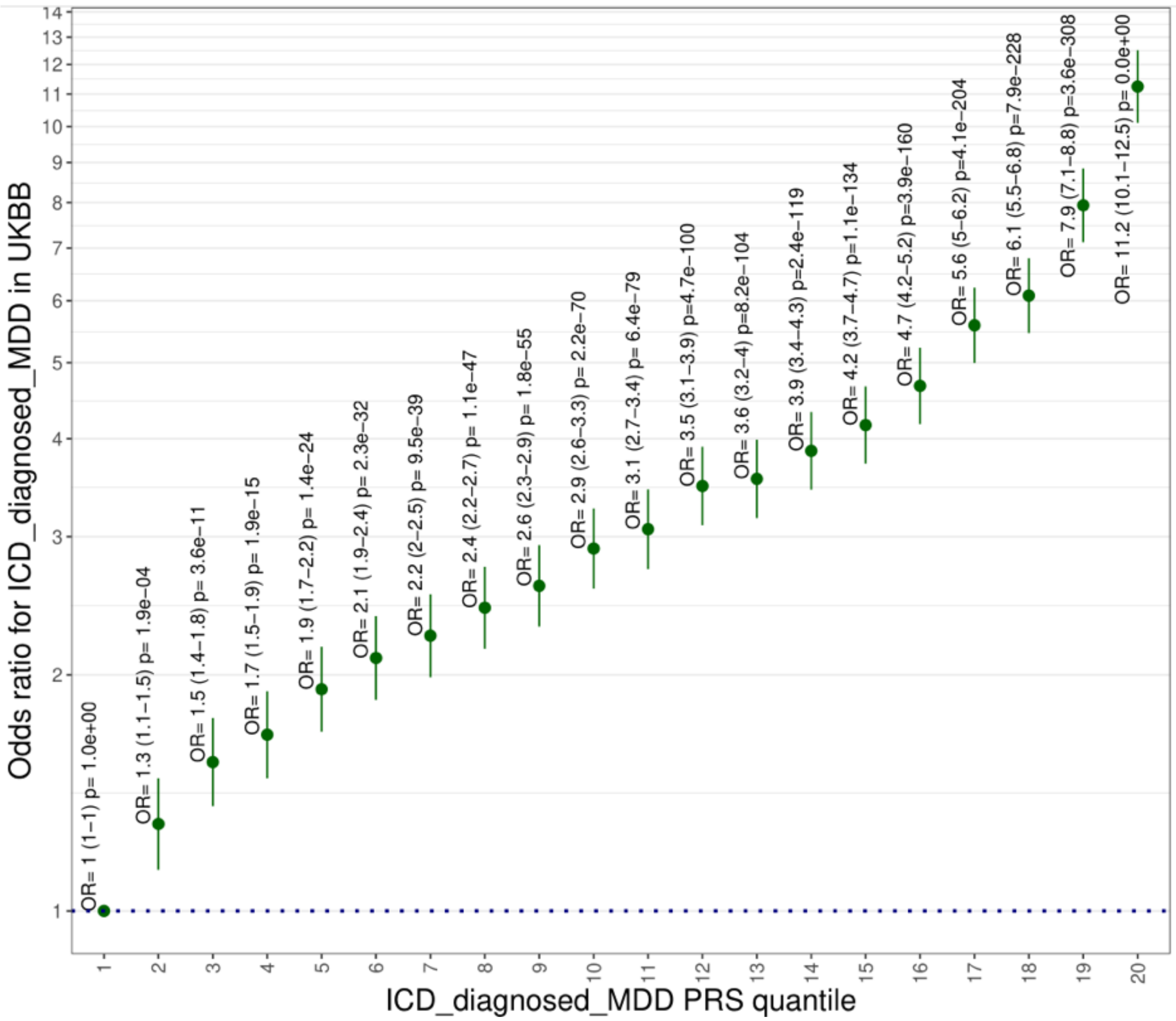

*Supplementary Figure 5: OR for MDD by Rare Variant IOGC Quantiles*

***The quantile plot shows the odds ratios and 95% confidence intervals for MDD for 20 quantiles of IOGC rare variant burden scores, from 1 (reference, and lowest PRS) to 20, in UKBB.***

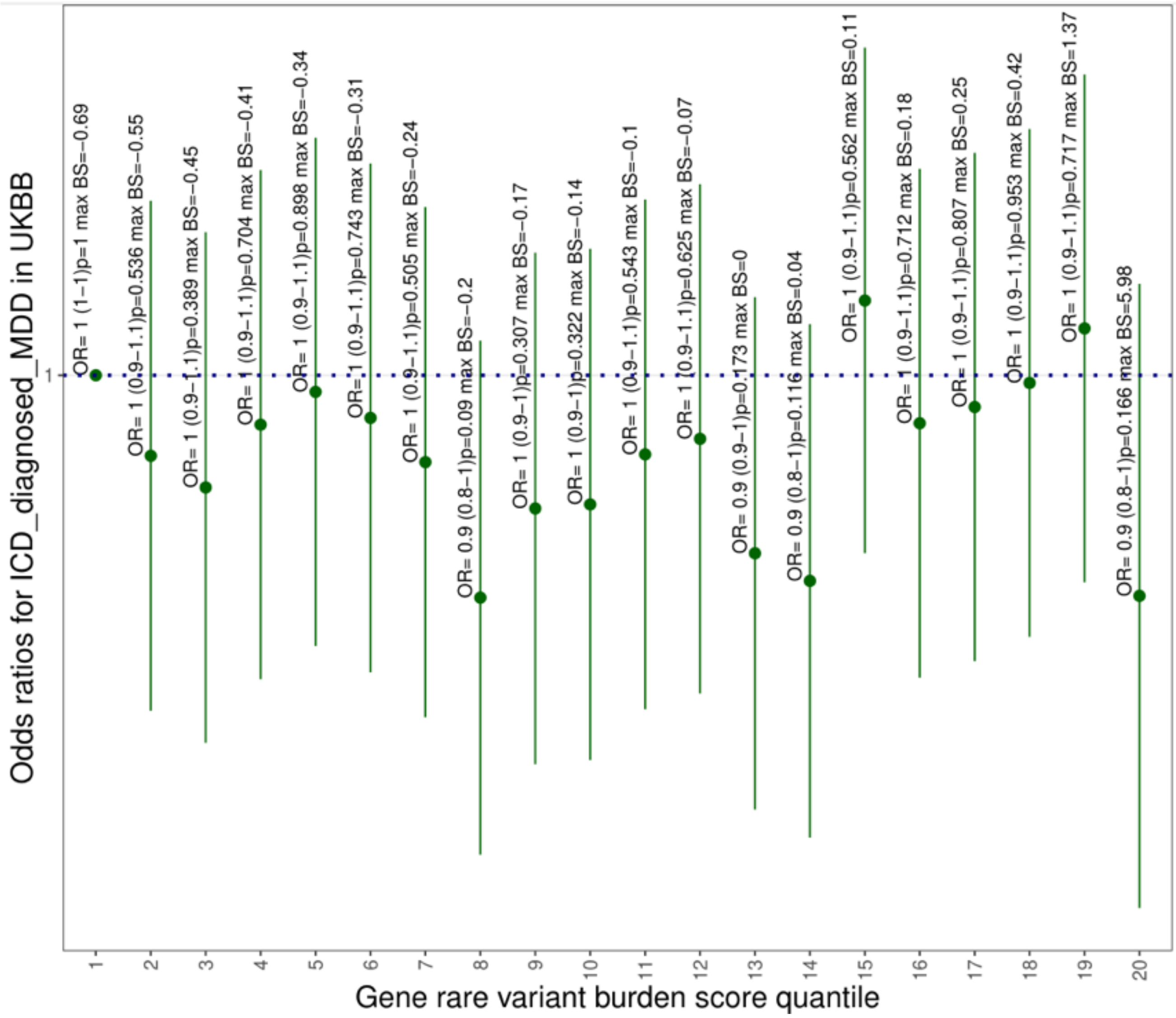

*Supplementary Figure 6: Positive Predictive Value for MDD by common-variant PRS and by high/low IOGC*

*The figure shows positive predictive values (PPVs) for MDD across pre-specified PRS centile thresholds (whole sample, <15th, <25th, <50th, <75th, >95th) separately within low- vs high-IOGC strata (dichotomised at the pre-specified IOGC quantile cutoff): in blue, low IOGC; in red, high IOGC.*

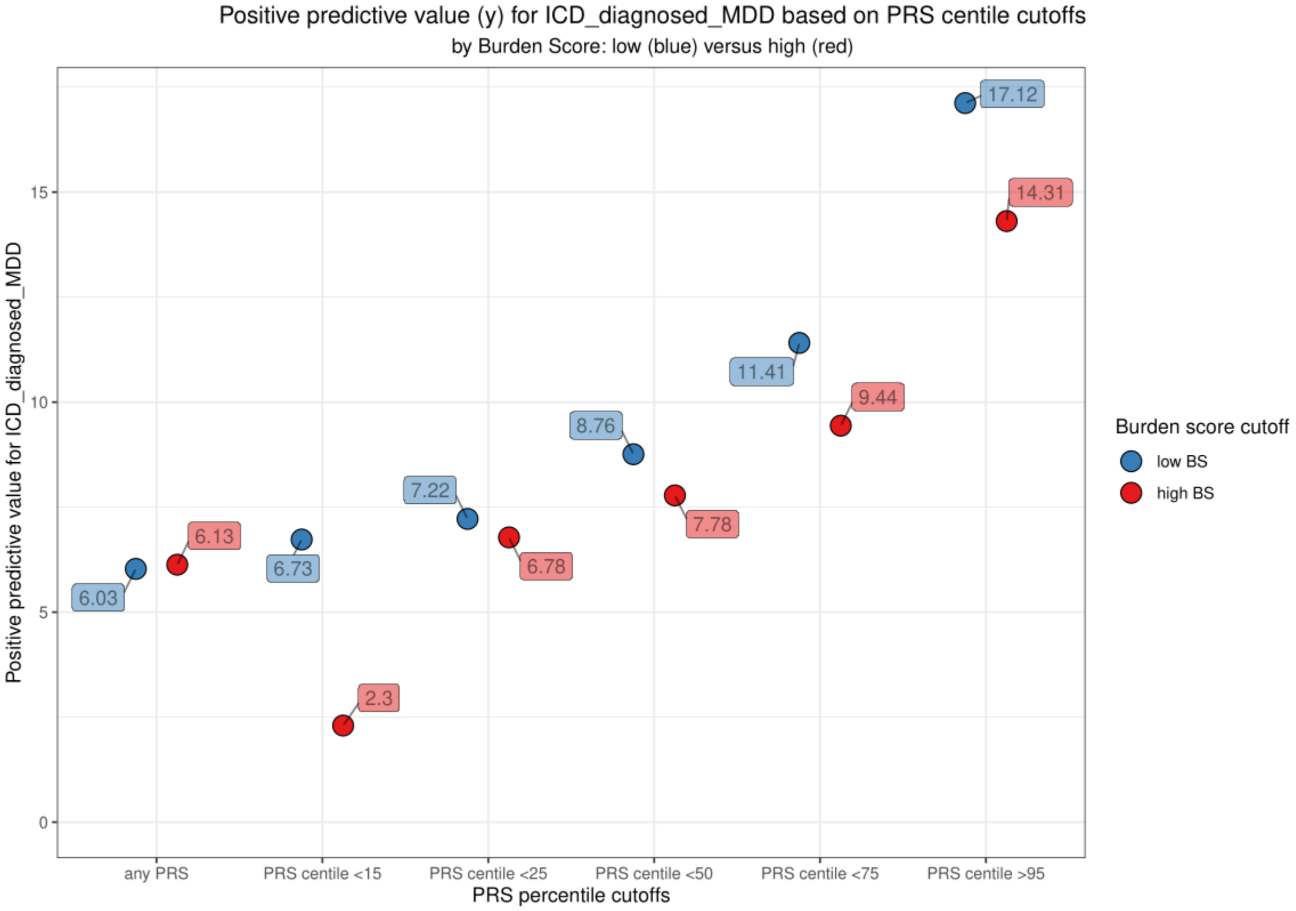

*Supplementary Figure 7: OR for HCM by Common Variant HCM PRS Quantiles*

*The quantile plot shows the odds ratios and 95% confidence intervals for HCM for 20 quantiles of original GWAS PRS, from 1 (reference, and lowest PRS) to 20, in UKBB.*

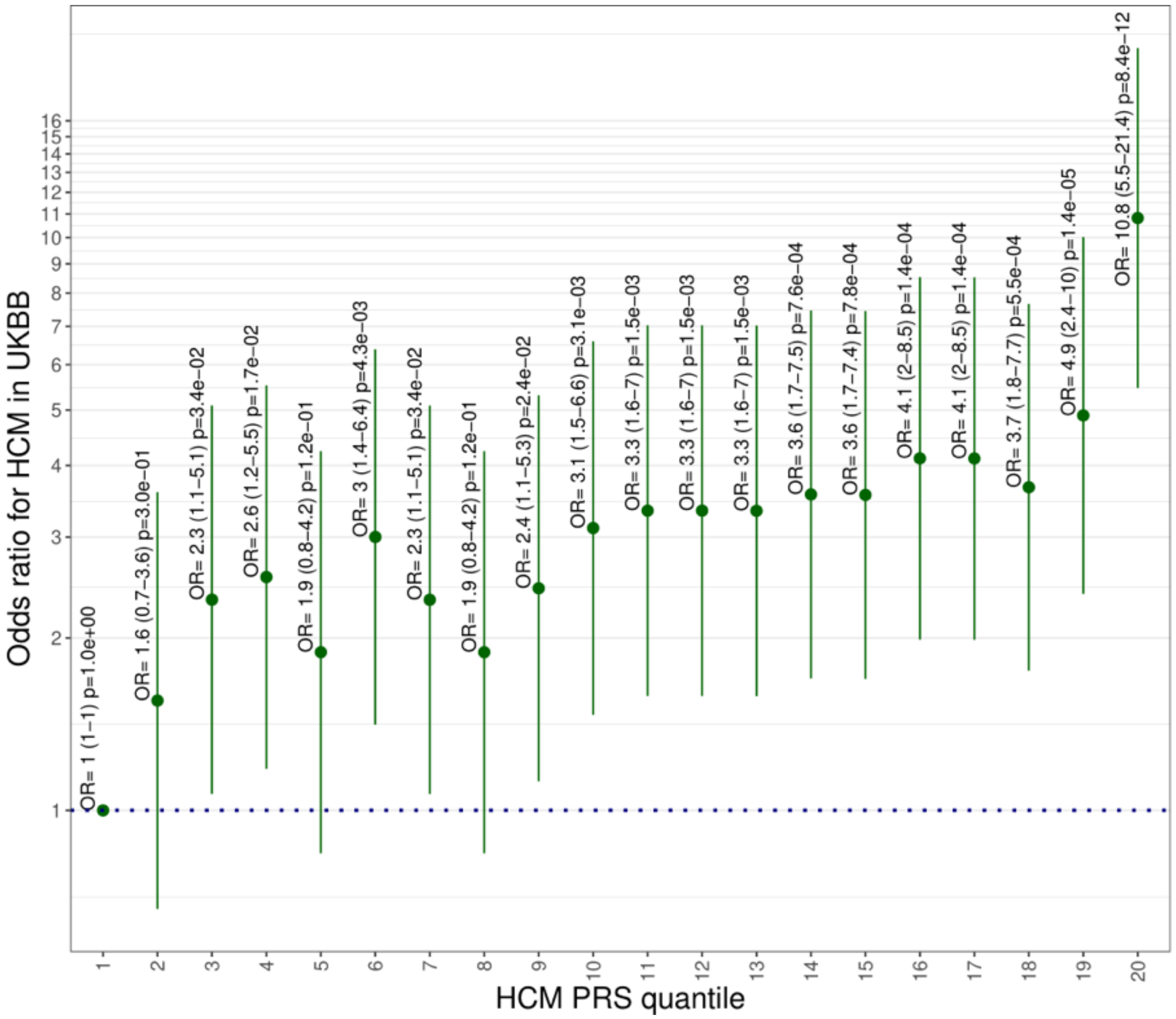

*Supplementary Figure 8: OR for HCM by Rare Variant IOGC Quantiles*

***The quantile plot shows the odds ratios and 95% confidence intervals for HCM for 20 quantiles of IOGC rare variant burden scores, from 1 (reference, and lowest PRS) to 20, in UKBB.***

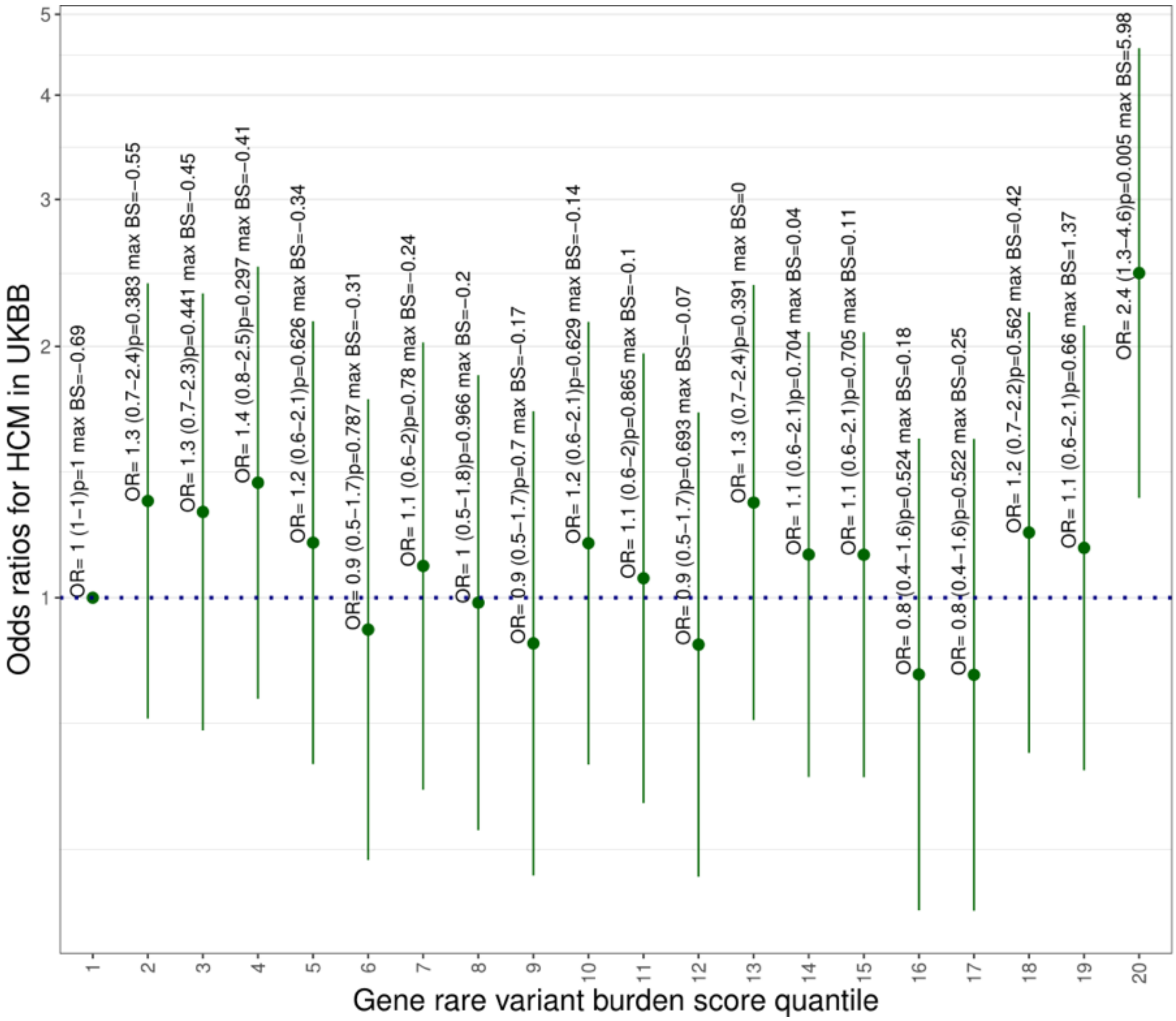

### *Supplementary Figure 9: Positive Predictive Value for HCM by common-variant PRS and by high/low IOGC*

*The figure shows positive predictive values (PPVs) for HCM across pre-specified PRS centile thresholds (whole sample, <15th, <25th, <50th, <75th, >95th) separately within low- vs high-IOGC strata (dichotomised at the pre-specified IOGC quantile cutoff): in blue, low IOGC; in red, high IOGC.*

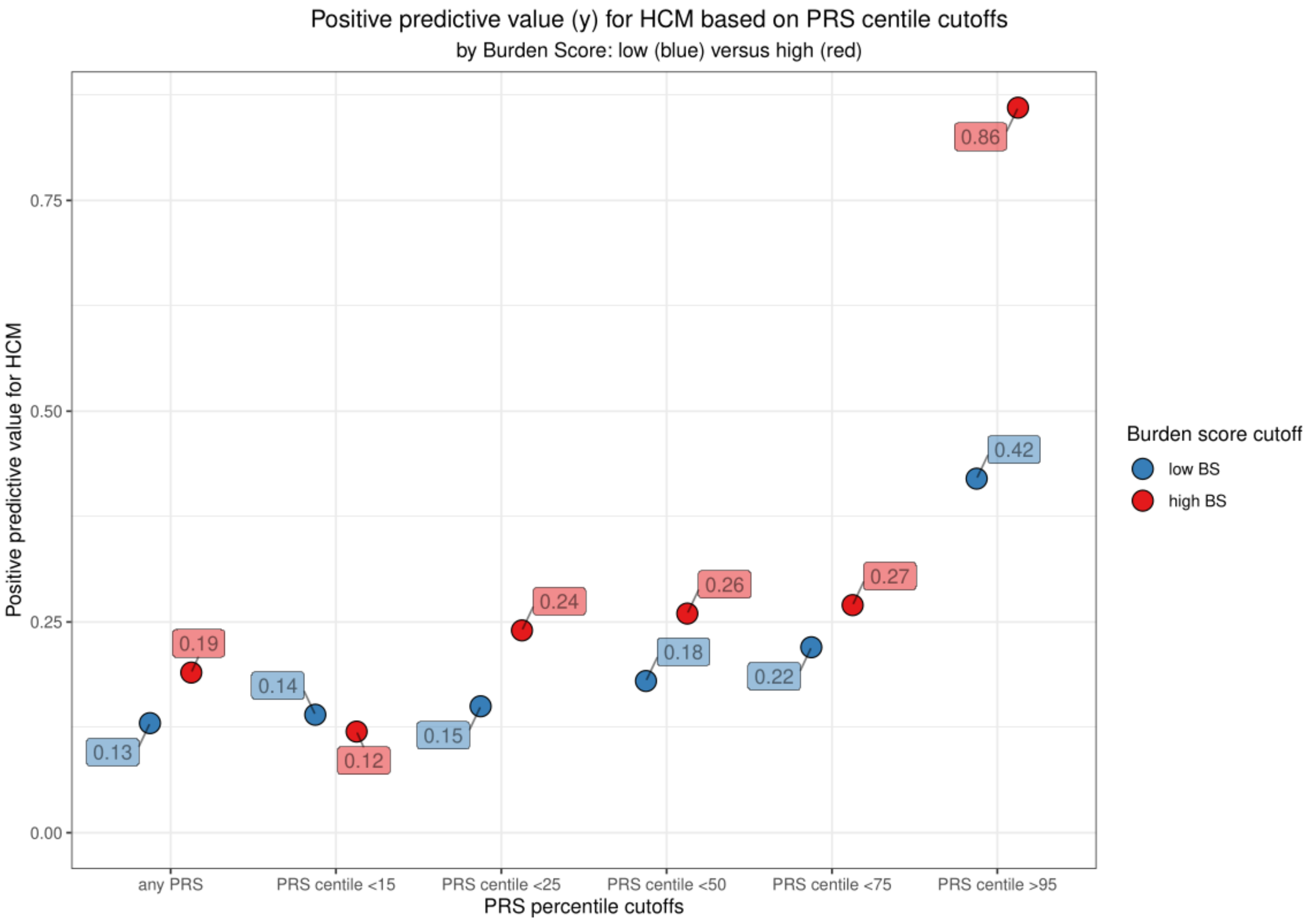

*Supplementary Figure 10: Positive Predictive Value for schizophrenia by common-variant PRS and by high/low randomised IOGC*

*The figure shows positive predictive values (PPVs) for schizophrenia across pre-specified PRS centile thresholds (whole sample, <15th, <25th, <50th, <75th, >95th) separately within low-vs high-* ***randomised*** *IOGC strata (dichotomised at the pre-specified IOGC quantile cutoff): in blue, low IOGC; in red, high IOGC.*

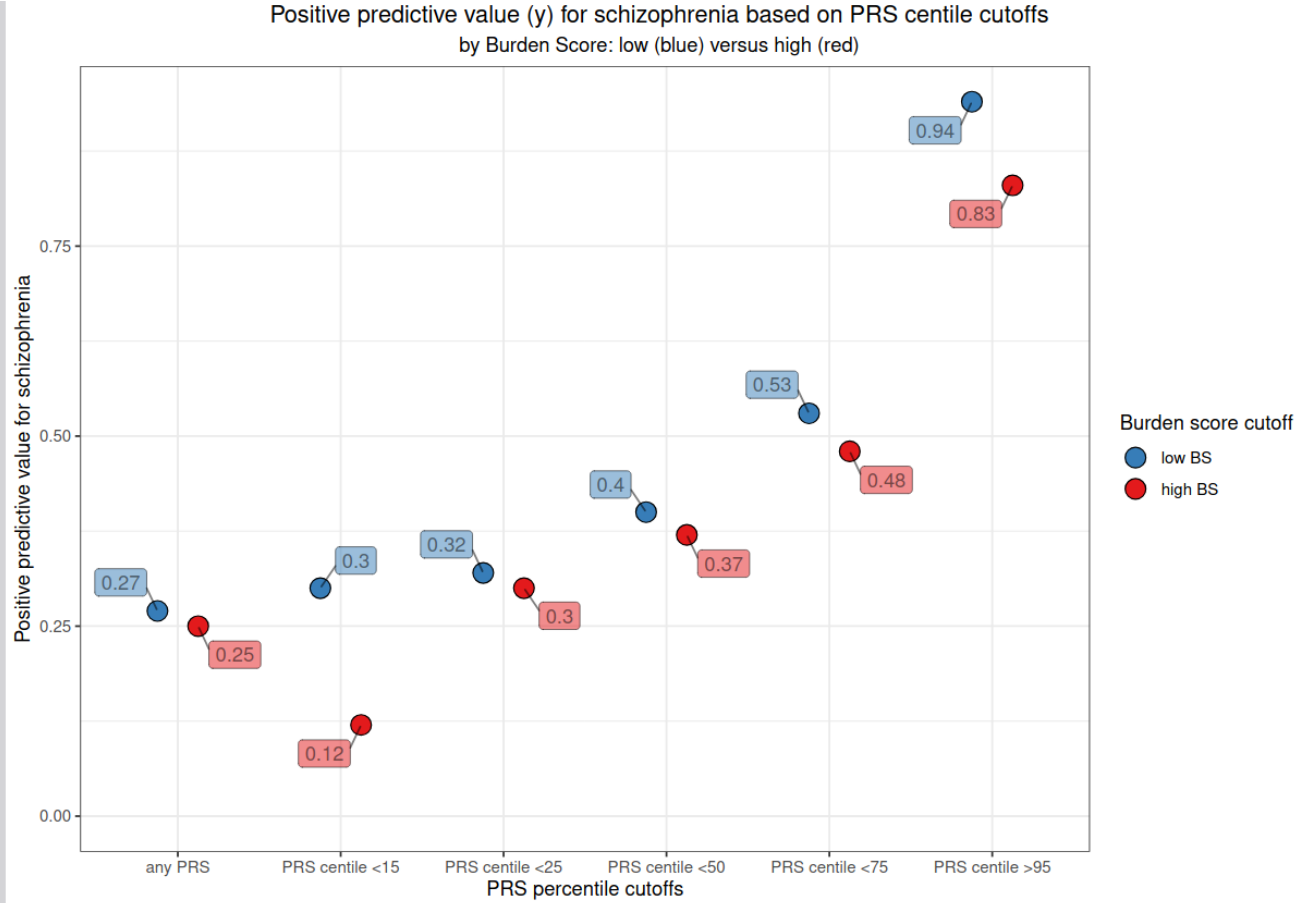

### *Supplementary Figure 11: Positive Predictive Value for HCM by common-variant PRS and by high/low randomised IOGC*

*The figure shows positive predictive values (PPVs) for HCM across pre-specified PRS centile thresholds (whole sample, <15th, <25th, <50th, <75th, >95th) separately within low- vs high-* ***randomised*** *IOGC strata (dichotomised at the pre-specified IOGC quantile cutoff): in blue, low IOGC; in red, high IOGC.*

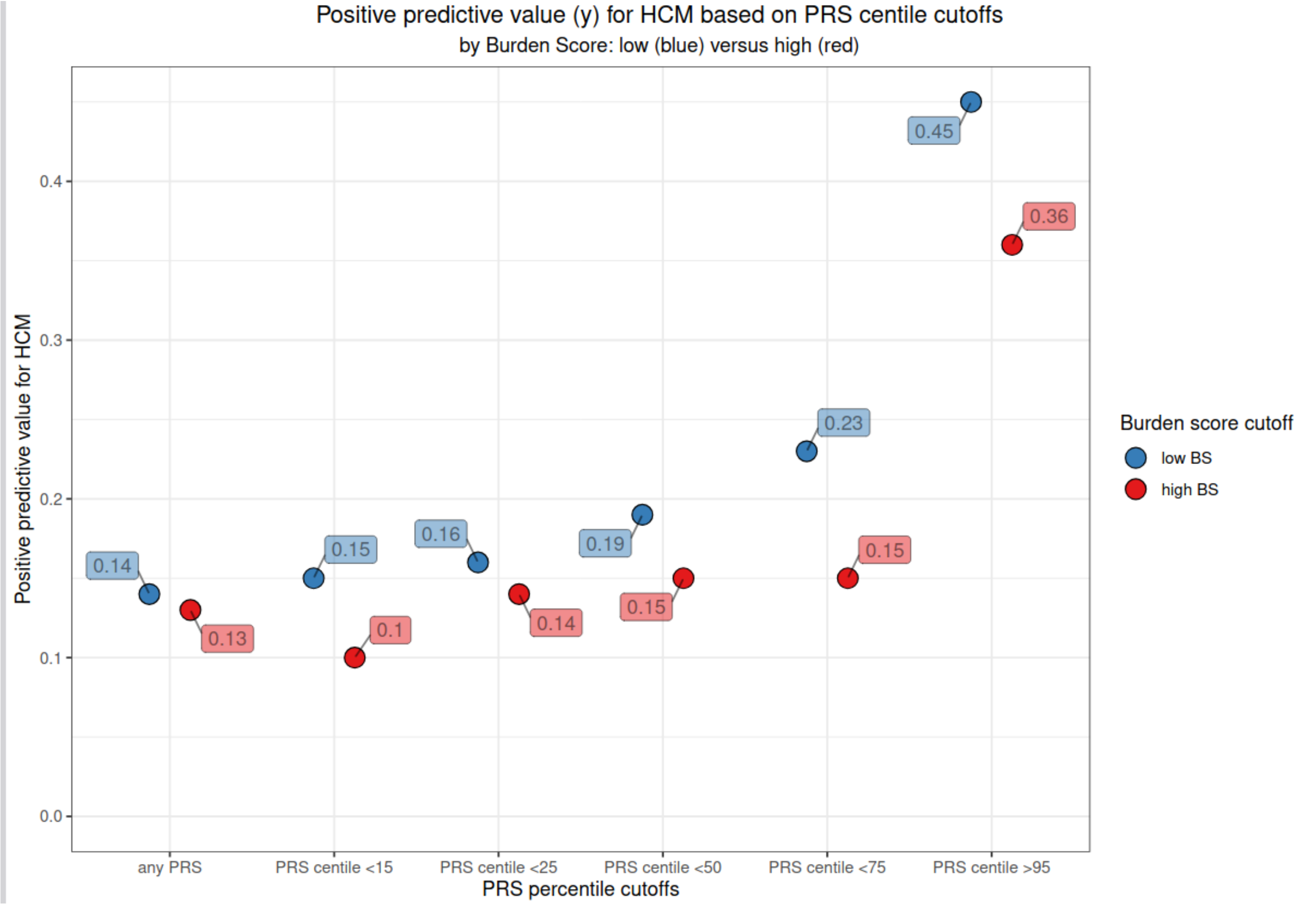
